# Perception of aversive stimuli of different gustatory modalities in an haematophagous insect, *Rhodnius prolixus*

**DOI:** 10.1101/616615

**Authors:** Santiago Masagué, Agustina Cano, Yamila Asparch, Romina B. Barrozo, Sebastian Minoli

## Abstract

Sensory aversion is an essential link for avoiding potential dangers. Here, we studied the chemical perception of aversive compounds of different gustatory modalities (salty and bitter) in the haematophagous kissing bug, *Rhodnius prolixus*. Over a walking arena, insects preferred a substrate embedded with 0.3 M NaCl or KCl rather than with distilled water. Same salts were avoided when prepared at 1 M. When NaCl and KCl were confronted, no preferences were evinced by insects. A pre-exposure to amiloride interfered with the repellency of NaCl and KCl equally, suggesting that amiloride-sensitive receptors are involved in the detection of both salts. Discriminative experiments were then performed to determine if *R. prolixus* can distinguish between these salts. An aversive operant conditioning involving either NaCl or KCl modulated the repellency of the conditioned salt, but also of the novel salt. A chemical pre-exposure to the salts did not to modify their repellency levels. When we crossed gustatory modalities by confronting NaCl to caffeine (*i.e.* a bitter stimulus) no innate preferences were evinced. Aversive operant conditionings with either NaCl or Caf rendered unspecific changes in the repellency of both compounds. A chemical pre-exposure to Caf modulated the response to Caf but not to NaCl, suggesting the existence of two independent neural pathways for the detection of salts and bitter compounds. Overall results suggest that *R. prolixus* cannot distinguish between NaCl and KCl but can distinguish between NaCl and Caf and generalizes the response between these two aversive stimuli of different gustatory modality.

**Summary statement:** Kissing-bugs use contact chemo-perception to avoid aversive substrates. They can sensory distinguish between salty (sodium chloride) and bitter (caffeine) tastes, but not between different salts (sodium and potassium chloride).

## 1. Introduction

Innate aversion is a highly adaptive behaviour that plays a key role in the survival of animals, as it can prevent them from suffering critical damages. Aversive stimuli can promote avoidance or escape in an individual by directly provoking pain (*e.g.* electric shock) or by alerting of the presence of a potential danger (*e.g.* alarm pheromone). In both cases, the peripheral sensory system is in charge of communicating the presence of a danger to the brain. Among the different sensory modalities, gustatory aversiveness acquires a high relevance, as it can help in preventing the ingestion of large quantities of toxic or harmful compounds that might be present in food. Taste is a quite simple system that allows the detection and differentiation of relative low number of stimuli as compared with the olfactory sense. For example, animals can detect more than thousands of different olfactory molecules, but they can only differentiate a dozen or less flavours (Liman *et al*., 2014; McGann, 2017).

In insects, taste or contact chemoreception sensilla can be found assembled in specific appendages such as proboscis, antennae or legs, but they can also be present all along the insect’s body (*e.g.* abdomen, ovipositor or wings). Even if they are called “taste” sensilla, they can bear functions other than those involved in food recognition, such as identification of suitable/unsuitable oviposition sites, detection of toxic substrates, recognition of congeners and sexual partners, among other. In any case, the functional unity of detection is the taste sensillum, which bears one apical pore and a few neurons inside. Typically, one of the neurons is a mechano-receptor that helps the animal to determine if the sensillum is contacting a surface or not, while the other/s is/are chemo-receptors (Singh, 1997). This arrangement can, however, change among species. Differently from the olfactory system in which each neuron possesses one only type of receptor and hence it’s specific for one odour, many gustatory receptor (GR) types can be present in each gustatory receptor neuron (GRN) (Freeman and Dahanukar, 2015). This is probably one of the reasons why only few tastants can be discriminated by mammals and insects (Masek and Scott, 2010; Spector and Kopka, 2002; Asparch *et al*., 2016). Even if this could be seen as a disadvantage, it might rather be an advantage for an animal, as it allows to group chemical stimuli that communicate the same signal (for example “toxic”) even if they have quite different molecular structures. For example, plants defence against herbivory includes the production of numerous toxic compounds that have a bitter taste (at least for humans) and generate aversive responses in animals. So even without being able to discern if two bitter molecules are the same or not, it would be useful for an animal simply to reject both of them. Contrarily, there are also cases in which determining the chemical identity of a tastant can make the difference between living or not. For example many alkaloids produced by plants to avoid being eaten have different toxicity levels, for what if an animal’s taste system could distinguish between them, in low-food availability conditions it might be very adaptive to feed over plants producing the least toxic ones (Glendinning, 1994; Glendinning *et al*., 2001; Ayestaran *et al*., 2010; Hurst *et al*., 2014). In haematophagous insects such as mosquitoes and kissing bugs, feeding responses were partially or even completely inhibited by the addition of alkaloids to appetitive artificial diets (Kessler *et al*., 2014; Pontes *et al*., 2014). Moreover, a walking substrate impregnated with two different alkaloids (caffeine and quinine) was repellent for *R. prolixus* (Asparch *et al*., 2016). Although the avoidance response generated by caffeine and quinine was quite similar, conditioning protocols allowed authors to determine that these insects are able to discriminate among them.

Besides avoiding potentially toxic compounds, animals need to ingest the correct amount of salts to maintain the ionic balance inside their bodies. Excessive ingestion of salts can have as much harmful consequences as the absence of salts in the diet. Therefore, the detection of salts and their concentrations in a potential food source becomes relevant for most animals, reason why it normally guides the decision making about feeding or not on a particular diet. In this search for homeostasis, sodium chloride (NaCl) seems to be a main actor. In blood feeders such as mosquitoes, kissing bugs, fleas and tse-tse flies, NaCl is crucial to elicit gorging under artificial feeding conditions (Friend and Smith, 1977). However, feeding rejection occurs if salt concentration is below or above vertebrate’s blood level (Pontes *et al*., 2017; Galun *et al*., 1963). In *R. prolixus*, NaCl detection seems to be driven be amiloride-sensitive receptors present on the GRNs of the pharyngeal sensilla (Pontes *et al*., 2017). However, no information is available about the effects that the detection of salts over the walking substrate might have on the behaviour of such haematophagous insects.

This work aims to disentangle the capacity of a disease vector insect, the haematophagous *R. prolixus*, to assess the gustatory quality of a walking substrate. We firstly explored if they are capable of perceiving different concentrations of two salts: NaCl and KCl. Then, we investigated if they are able to distinguish between these two salts (*i.e.* two stimuli from the same gustatory modality) or between NaCl and a bitter compound as caffeine (*i.e.* two stimuli from different gustatory modality). Different experimental approaches were applied to analyse the discriminatory capacities of *R. prolixus*. Along this work we assume that if a particular treatment modulates the response to one stimulus but not to the other, insects are capable of discriminating among them.

## 2. Materials and methods

### 2.1. Insects

*Rhodnius prolixus* (Heteroptera: Reduviidae: Triatominae) was reared in the laboratory at constant 28 ± 1 °C, 60 ± 20 % relative humidity and 12 : 12 h. L / D photoperiod cycle. Recently moulted 5^th^ instar nymphs were kept unfed for 14 days and randomly assigned to the different treatments along this work. Each nymph was used only in one experiment and then returned to the rearing chamber. All animals were handled according to the biosafety rules of the Hygiene and Safety Service of the University of Buenos Aires.

Being nocturnal insects, all experiments were carried out during the first hours of the insects’ scotophase (*i.e.* 1 - 6 h. after lights were turned-off) in a dark experimental room. The temperature of the room was set to 25 ± 1 °C for all assays. This spatio-temporal arrangement allowed us to exclude external visual cues during experiments and at the same time match the maximal activity period described for triatomines (Lazzari, 1992).

### 2.2. Stimuli

Sodium chloride (NaCl), potassium chloride (KCl) and caffeine anhydrous (Caf) were purchased from Biopack (Buenos Aires, Argentine). Amiloride hydrochloride hydrate was purchased in Sigma-Aldrich® (St. Louis, U.S.A). The different stimulus solutions (0.3 M and 1 M NaCl; 0.3 M and 1 M KCl; 0.1 M Caf; 1 mM amiloride) were prepared in distilled water, stored at 4°C and used in the same week.

### 2.3. Taste preference assay: two-choice walking arena

Individual assays were performed in a rectangular acrylic box (10 × 5 cm) whose floor was completely covered with a filter paper (see upper panel of all figures). A line was drawn on the paper at the centre of the arena to define two equal square zones (5 × 5 cm). Over the paper of each zone 100 µl of a stimulus solution (*i.e.* distilled water, 0.3 or 1M NaCl, 0.3 or 1M KCl or 0.1M Caf) were homogeneously spread using a micropipette. In this way, according to the experimental series, the following chemical two-choices were generated: H_2_O/NaCl, H_2_O/KCl, KCl/NaCl, H_2_O/Caf or Caf/NaCl. After one minute, an insect was gently located at the centre of the arena and covered with an inverted flask for another 1 minute familiarization with the experimental setup. Then, the flask was gently removed and the experiment started. The position of the freely-walking insect in relation to the position of the added stimuli was continuously recorded during 4 minutes using an infrared-sensitive video-camera connected to a digital recorder. Note that the investigator and the recording system (except the camera) were located outside the experimental room to avoid the addition of potential cues that could be detected by the insects.

From the video recordings, the time spent in each side of the arena was computed for each insect. A Preference Index (PI) was calculated as the time spent at the stimulus “A” side (*St A*), minus the time spent at the stimulus “B” side (*St B*), divided by the total experimental time:

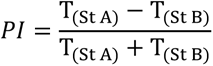

PIs close to −1 or 1 show preference for “B” or “A”, respectively. PIs near 0 indicate lack of preference. Thirty replicates were performed for each experimental series. Insects were used once and then discarded. In each experimental series, the stimulus was assigned to the “left” or “right” side of the arena in a pseudorandom manner.

### 2.4. Discriminative experiments: pre-treatments

Discriminative experiments were performed to discern if insects can distinguish between different aversive stimuli. Different pre-treatments were applied before the taste preference assays.

#### 2.4.1. Pre-exposure to amiloride

Amiloride is a pyrazine known to block the detection of NaCl in animals by interfering with the functioning of epithelial sodium channels (ENaC) (Kellengerger and Schild, 2002). We pre-expose insects to amiloride to examine if there is a modulation of the perception of NaCl, KCl or both salts.

Insects were individually exposed to amiloride inside a plastic vial (2 cm diam., 3 cm height, see upper panel of Fig. 2) whose floor was covered with a filter paper loaded with 100 µl of H_2_O (control series) or of 1 mM amiloride. One insect was gently placed over the paper inside the vial and left to freely walk during 1 minute. During this time, legs, antennae, proboscis and/or other parts of the body could contact the H_2_O or the amiloride. The insect was then removed, maintained in a different clean flask for 2 minutes and then transferred to the two-choice arena where its chemical preference was tested as explain in section 2.3.

**Figure 1.**
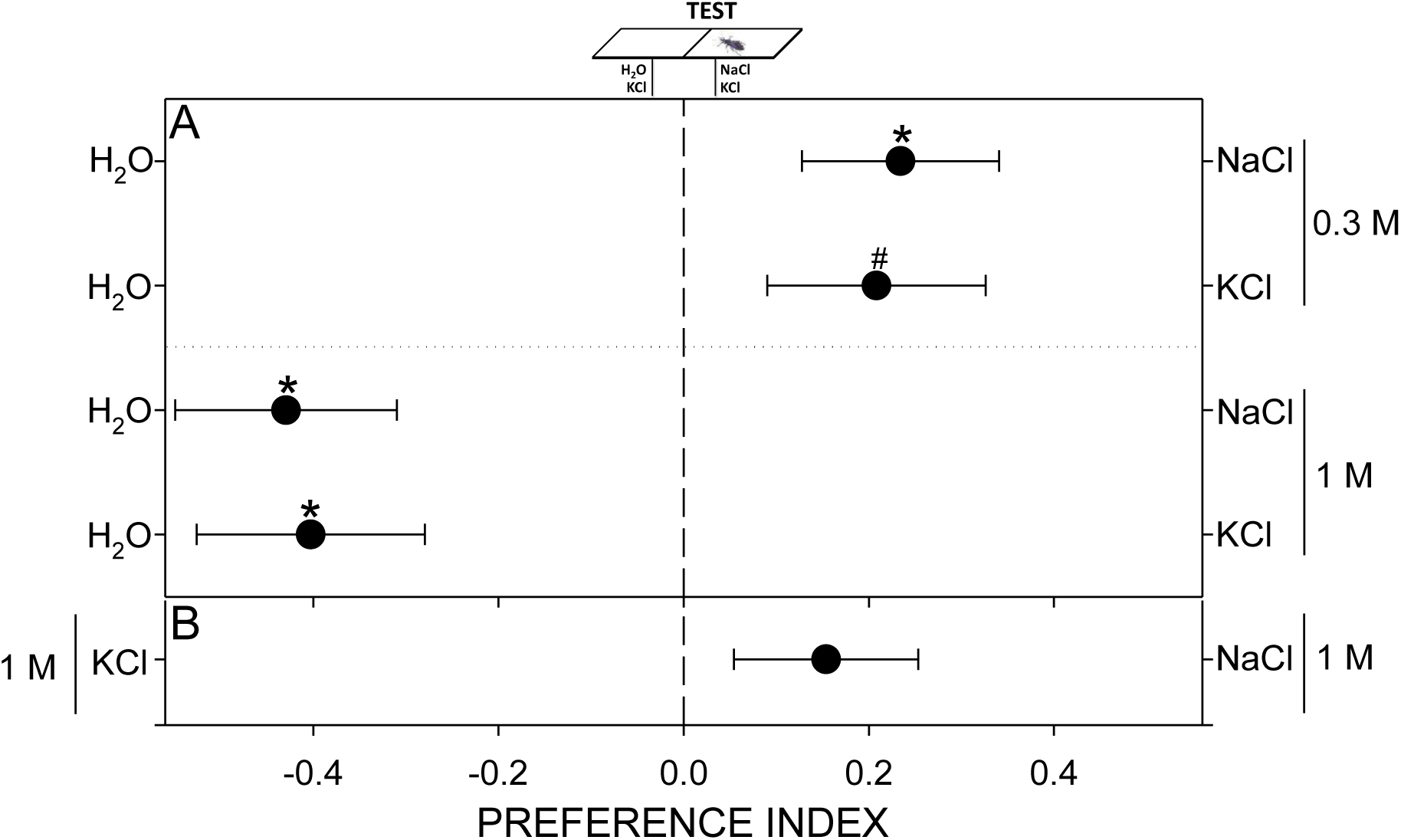
Concentration-dependent responses to NaCl and KCl. Insects preferred low (0.3 M) and avoided high (1 M) concentrations of both salts confronted to H_2_O (A). Insects exhibited no preference when both salts were confronted (B). The Preference Index expresses the relative time spent at each side of the arena: 0 = equal time at each side, −1 and 1 = full time spent at the left or right side of the arena, respectively. Each point represents the mean (± s.e.m.) of 30 replicates. Asterisks denote statistical differences (p < 0.05) after a One-Sample T-Test with expected value = 0. Numeral shows a p-value of 0.056.

**Figure 2.**
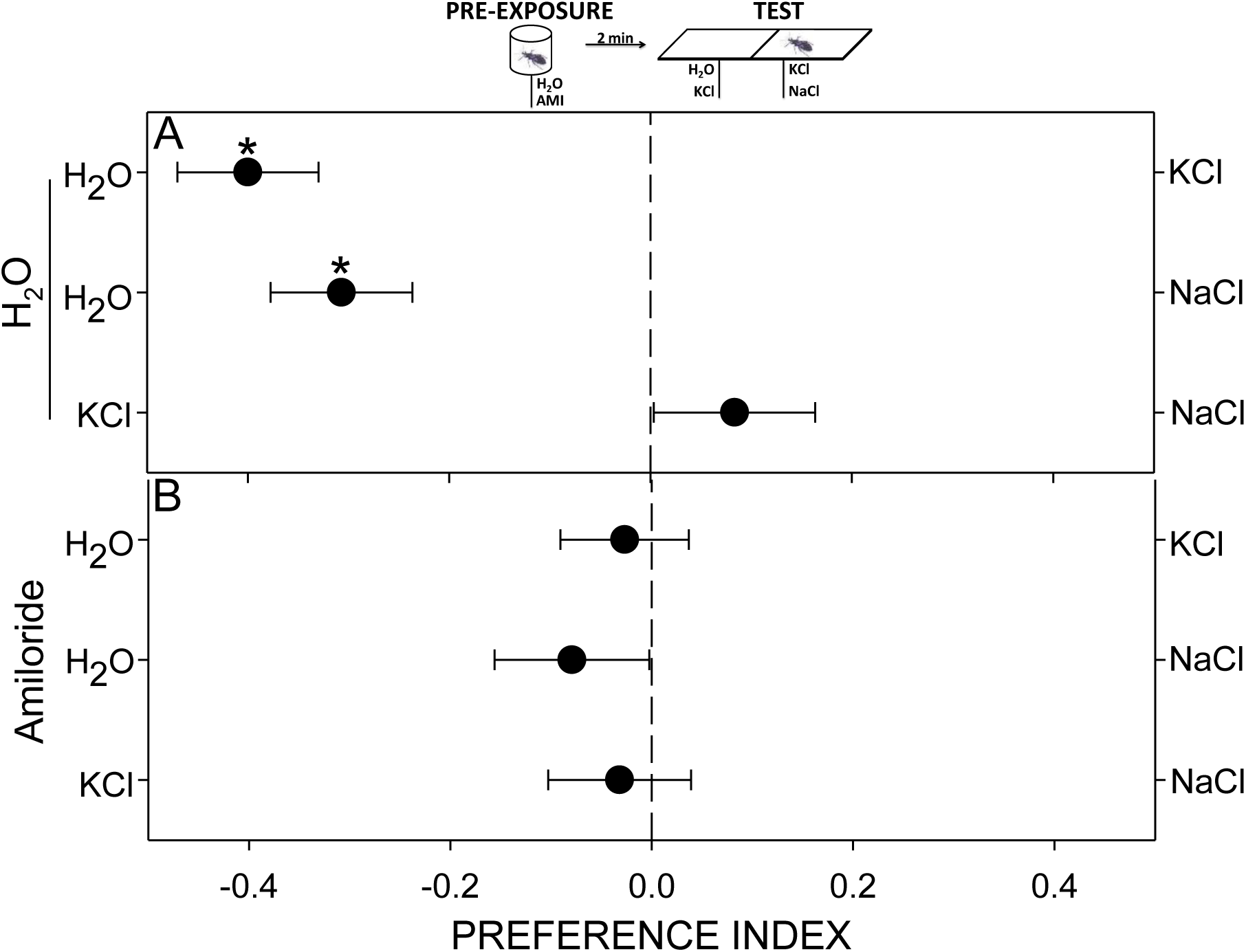
Amiloride blocks salt perception. A pre-exposure to H_2_O did not modify the innate repellence of insects to both salts (A). A pre-exposure to amiloride interfered with the perception of both salts (B). The Preference Index expresses the relative time spent at each side of the arena: 0 = equal time at each side, −1 and 1 = full time spent at the left or right side of the arena, respectively. Each point represents the mean (± s.e.m.) of 30 replicates. Asterisks denote statistical differences (p < 0.05) after a One-Sample T-Test with expected value = 0.

#### 2.4.2. Discriminative learning: operant aversive conditioning

Associative learning protocols are widely applied in discriminatory studies, as they can help in determining if an individual can detect as different two stimuli generating the same behavioural output. It is assumed that if after a conditioning protocol involving stimulus “A” there is a specific modulation of the response to “A” and not to “B”, the individual can distinguish between “A” and “B”.

An operant aversive conditioning of 4 minutes was individually applied to the insects over the experimental arena described in section 2.3. But, to deliver a negative reinforcement to the insects, a vortex-type laboratory mixer was mechanically attached to the arena (see upper panel of Figs. 3, 4, 7 and S1), allowing us to voluntarily deliver a vibration disturbance via an external switch. For each conditioning, 100 µl of the different compounds were loaded at each side of the arena as explained before (see section 2.3), *i.e.* KCl/NaCl, H_2_O/NaCl, H_2_O/KCl or Caf/NaCl. One of the compounds was predefined as the punished stimulus, the other as the safe one (note that in the text and the figures a negative sign (-) is added beside the stimulus punished during training). An insect was then located at the centre of the arena, covered with an inverted flask for one minute for familiarization with the experimental setup. Then the flask was gently removed, and the aversive conditioning started. During the following 4 minutes of training, the insect could control the occurrence of the mechanical disturbance by its position: every time it entered the side of the arena loaded with the punished stimulus, the vortex mixer was switched on, generating a mechanical disturbance for the insects. Conversely, the vibration stopped whenever the insect entered the safe side of the arena.

Unpaired yoke control series were carried out to verify the associative origin of the behavioural modulation. For this, during 4 minutes each individual received a negative reinforcement independently from its position in the experimental arena (note that in the text and the figures negative signs (-) are added beside both stimuli during training). To match the intensity of the vibration received by insects of the conditioning groups, the timing, frequency and duration of the vibration delivered to yoke control insects were copied from the previously conditioned insect. In this way, individuals of these series received the same amount of vibration than those from the experimental series, but in this case dissociated from the position of the chemical stimuli.

In all cases, the behaviour of each individual during the conditioning time was registered in video as explained before, and the individual PIs were computed. Once the conditioning ended, the insect was removed, put in an individual flask for 2 minutes and then transferred to the two-choice arena where its chemical preference (*i.e.* without vibrations delivered) was tested as explain in section 2.3. Note that for these series, separated PIs are registered for training (triangles in the figures) and test phases (circles in the figures).

#### 2.4.3. Compound-specific chemical pre-exposure

Long chemical exposures generally give rise to reduced responsiveness in animals. Two possible processes might be responsible for this modulation: 1- a peripheral sensory adaptation, in which receptors stop sending electrical information to the brain and consequently the behavioural response decreases, 2- an habituation, in which peripheral receptors send the electrical report to the brain but this information is processed and the response also decreases. In any case, it is assumed that if after an exposure to stimulus “A” there is a specific modulation of the response to “A” and not to “B”, the individual can distinguish peripherally between “A” and “B”.

A non-associative pre-exposure to NaCl, KCl or Caf was applied individually to the insects before testing their chemical preferences. Pre-exposure procedure was achieved in a plastic vial (2 cm diam., 3 cm height, see upper panel of Figs. 5, 8) whose floor was covered with a filter paper loaded with either 100 µl of 1 M NaCl, of 1 M KCl or of 0.1 M Caf. In control series the paper was loaded with 100 µl of H_2_O. One insect was gently placed inside the vial over the paper and left to freely walk during 60 minutes. During this time, legs, antennae, proboscis and/or other parts of the body could contact the chemical stimuli. Once the pre-exposure ended, the insect was removed, put in an individual flask for 2 minutes, and then its chemical preference was tested as explained in section 2.3.

### 2.5. Data and analyses

Thirty insects were tested in each experimental series. One PI was computed for each insect. The mean PI of each series was statistically compared against the expected value if there were no chemical preferences, *i.e.* “0” by applying One-Sample T-Tests (α = 0.05). Normality and homoscedasticity were verified in all data series. All figures represent the mean Preference Index (x-axis) and the chemical compounds presented at each side of the arena (y-axis). Asterisks denote statistical differences between the PI and the value 0 (p < 0.05).

## 3. Results

### 3.1. Salt perception: concentration-dependent behaviour

When the different salts were confronted to distilled water, *R. prolixus’* responses were reliant on the tested concentrations. Insects preferred to walk over the 0.3 M NaCl solution rather than over water (Fig. 1A, H_2_O/NaCl, T = 2.2, p < 0.05). A similar trend was observed for 0.3 M KCl (H_2_O/KCl, T = 1.8, p = 0.056), although the PI was borderline not significantly different from 0. Conversely, bugs avoided a 1 M solution of both salts (H_2_O/NaCl, T = −3.6, p < 0.01; H_2_O/KCl, T = −3.3, p < 0.01).

These results show that *R. prolixus* can detect the presence of both salts over the walking substrate, and that the concentration perceived determines the type of response evinced by these bugs. In fact, concentrations close to those found over the skin of a potential host (*i.e.* around 0.1 M) were preferred, while higher ones, which could probably alter negatively the internal homeostasis of the insect, were repellent. However, regardless the concentration, and as the responses of insects to both salts were quite similar, next experimental series were designed to determine if these insects can detect both salts as different compounds or not. Experiments presented here onwards were carried out using the aversive concentration of NaCl and KCl, *i.e.* 1 M.

### 3.2 Detection of NaCl and KCl: different or same receptors?

#### 3.2.1. Innate response to the NaCl/KCl simultaneous presentation

As a first attempt to examine if these bugs can detect NaCl and KCl as different compounds, we offered them the possibility to choose between the aversive concentration of both salts (*i.e.* 1 M). We assumed that the expression of any spatial preference over the arena would indicate a differential perception of both stimuli. However, in this situation insects exhibited no preference for one or the other salt (Fig. 1B, T = 1.5, p > 0.05). This lack of preference exhibited could be the consequence of two quite opposite situations: 1- insects’ sensory system is unable to distinguish between both salts, or 2- insects can detect both salts as different compounds, but the triggered responses are similar in sign and intensity, *i.e.* NaCl and KCl are equally repellent. Next experimental series were performed to tackle this uncertainty.

#### 3.2.2. Inhibition of NaCl and KCl detection by amiloride

We tested here if amiloride can block the detection of salts in *R. prolixus* and if its action is specific for the detection of NaCl or if it interferes with the detection of KCl too. We assume that if the effect is specific for one salt, insects can distinguish between them.

Insects pre-exposed to H_2_O avoided both salts when faced to H_2_O (Fig. 2A, H_2_O/KCl, T= −5.7, p < 0.001; H_2_O/NaCl, T = −4.4, p < 0.001) and exhibited no preference when confronted to each other (KCl/NaCl, T = 1, p > 0.05). These control series show that there is no effect of pre-exposing insects to H_2_O. However, after a pre-exposure to amiloride, insects presented random distributions over the arena in all cases (Fig. 2B, H_2_O/KCl, T = −0.4, p > 0.05; H_2_O/NaCl, T = −1.0, p > 0.05; KCl/NaCl, T = 0.4, p > 0.05), evincing an unspecific lost in their capacity to perceive both salts.

Amiloride showed to efficiently inhibit salt detection in *R. prolixus*. The lack of specificity for NaCl or KCl suggests that the detection of both salts might be achieved by the same or at least by similar receptors, sensitive to amiloride blockage.

#### 3.2.3. Aversive conditioning with salts

Taking in consideration the results of sections 3.2.1 and 3.2.2, which show that NaCl and KCl generate the same avoidance response (see Figs. 1, 2), we applied operant aversive protocols to find out if *R. prolixus* can distinguish between these salts. We assume that the occurrence of a salt-specific conditioning indicates the existence of a differential detection pathway.

We firstly confronted KCl with NaCl during trainings, pairing the mechanical punishment to one of the salts, aiming to generate a specific aversive association during subsequent tests. Our results show that the mechanical vibration was indeed perceived as a negative reinforcement by bugs, as during conditionings insects avoided the punished side of the arena regardless if it contained KCl or NaCl (Fig. 3A, triangle KCl/NaCl(-), T = −8.0, p < 0.001; Fig. 3B, triangle KCl(-)/NaCl, T = −4.6, p < 0.001). However, in subsequent KCl/NaCl tests, insects exhibited no preference for one or the other salt, evincing that the conditioning did not affect their innate behaviour, (Fig. 3A, circle KCl/NaCl, T = 0.3, p > 0.05; Fig. 3B, circle KCl/NaCl, T = 0.4, p > 0.05).

**Figure 3.**
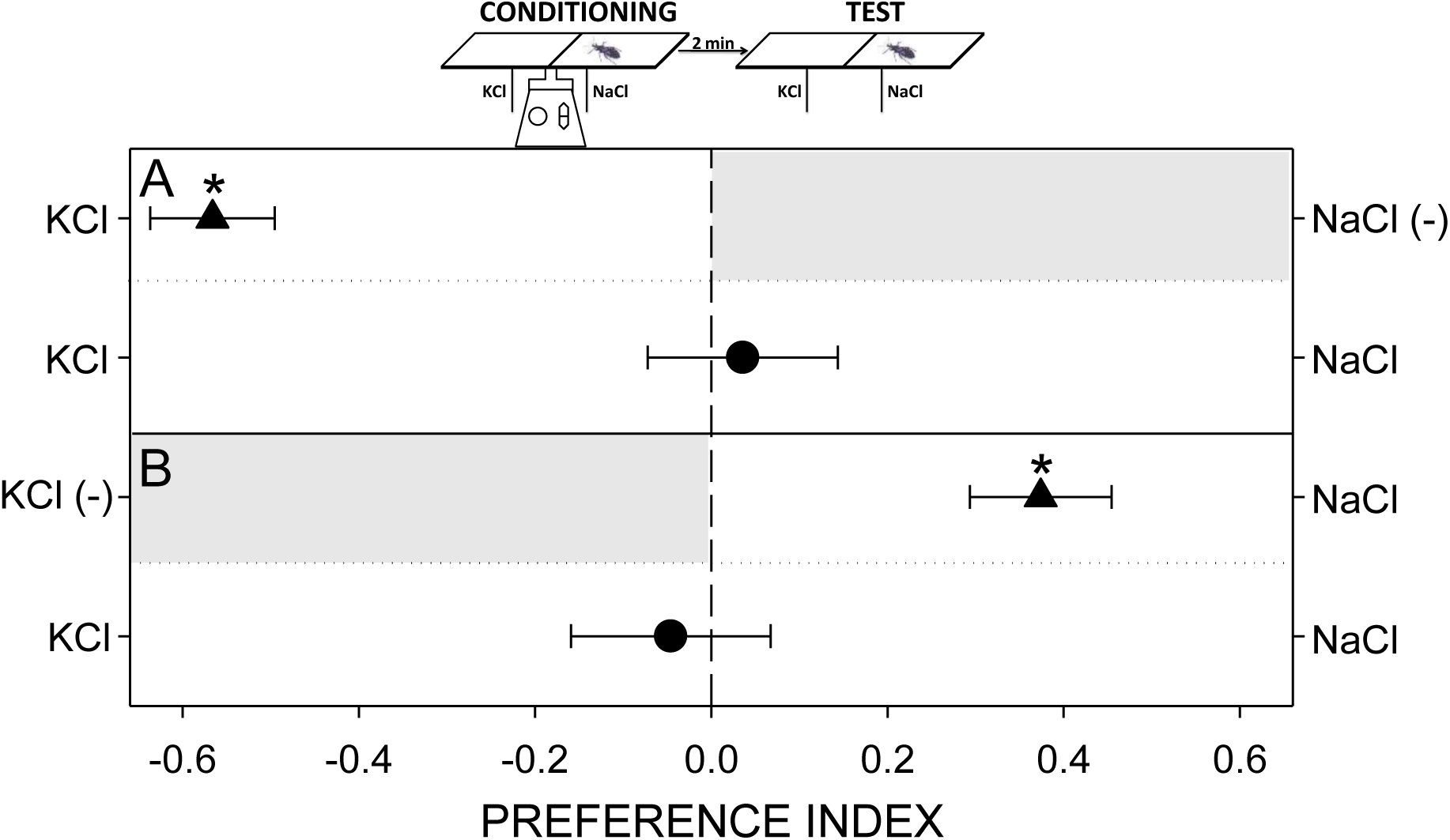
Salts discriminative assays through an operant conditioning protocol with NaCl(-) or KCl(-). During trainings (triangles in A and B) insects avoided the punished side of the arena (grey shadowed) regardless if it was loaded with NaCl or KCl. During tests (circles in A and B), no modifications of the innate behaviour were evinced, *i.e.* no preference when NaCl and KCl were confronted. The Preference Index expresses the relative time spent at each side of the arena: 0 = equal time at each side, −1 and 1 = full time spent at the left or right side of the arena, respectively. Each point represents the mean (± s.e.m.) of 30 replicates. Asterisks denote statistical differences (p < 0.05) after a One-Sample T-Test with expected value = 0.

Three different situations could explain this lack of preference during tests: 1- insects could be detecting both salts as different but the conditioning protocol could not be efficient (*e.g.* too short in duration, salience of the conditioned stimulus too low, negative reinforcement not enough, etc.), failing in modifying the innate lack of preference, 2- insects could be incapable of distinguishing between salts and then perceive an homogeneous substrate during both, training and test, or 3- insects could distinguish between salts and the conditioning protocol could be efficient, but they are generalizing what they have learnt for one salt to the other salt. Next series intend to separate and discard some of these options.

To determine if they are indeed capable of learning under our experimental conditions, we applied a similar conditioning protocol but involving only one salt and H_2_O during training. As we know from previous series that 1 M NaCl and KCl are aversive for these insects (see Fig. 1A), the negative reinforcement was applied at the H_2_O side of the arena. Once again, the mechanical vibration was efficient as negative reinforcement, even if insects had to remain in the salty side of the arena (Fig. 4A, triangle H_2_O(-)/NaCl, T = 5.2, p < 0.001; Fig. 4B, triangle H_2_O(-)/KCl, T = 5.4, p < 0.001). During tests insects continued to prefer the salty side of the arena rather than the H_2_O side (Fig. 4A, circle H_2_O/NaCl, T = 2.5, p < 0.05; Fig. 4B, circle H_2_O/KCl, T = 3.1, p < 0.01). Surprisingly, they also preferred the side of the arena containing a salt that was not presented during training (Fig. 4A, circle H_2_O/KCl, T = 3.2, p < 0.01; Fig. 4B, circle H_2_O/NaCl, T = 3.1, p < 0.01). Regardless the salt used during the aversive conditioning, no preferences were expressed in tests in which both salts were confronted (Fig. 4A, circle KCl/NaCl, T = 0.5, p > 0.05; Fig. 4B, circle KCl/NaCl, T = 0.5, p > 0.05).

**Figure 4.**
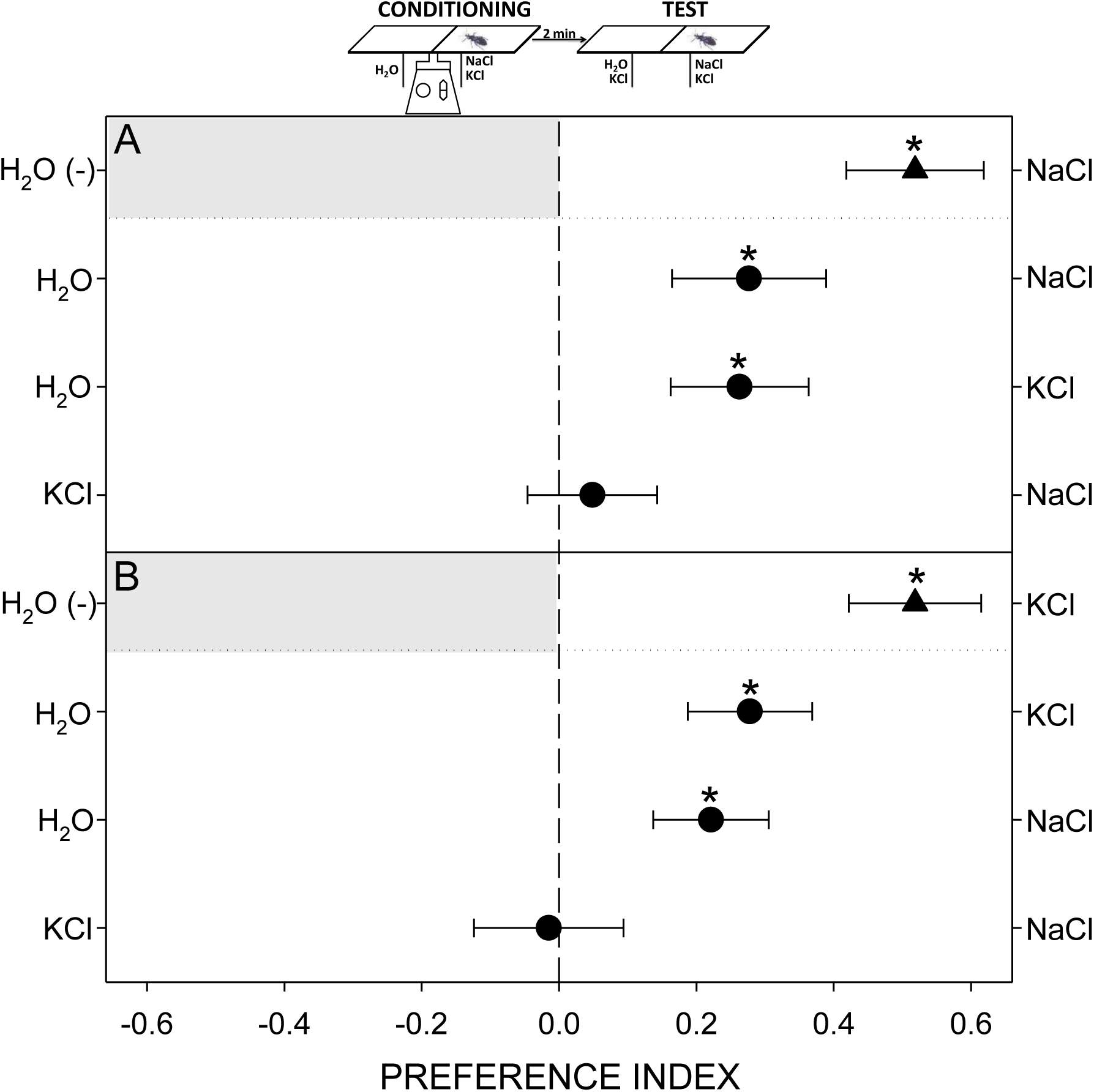
Salts discriminative assays through an operant conditioning protocol with H_2_O(-). During trainings (triangles in A and B) insects avoided the punished side of the arena (grey shadowed). During tests (circles in A and B), they preferred the salty side of the arena regardless if it contained the same salt used during training or the novel one. When confronted, no preference for one or the other salt was evinced. The Preference Index expresses the relative time spent at each side of the arena: 0 = equal time at each side, −1 and 1 = full time spent at the left or right side of the arena, respectively. Each point represents the mean (± s.e.m.) of 30 replicates. Asterisks denote statistical differences (p < 0.05) after a One-Sample T-Test with expected value = 0.

These results clearly show that *R. prolixus* can modify their chemical preferences after an aversive operant conditioning, *i.e.* they can learn to avoid punished stimuli. However, results evince that the observed experience-dependent plasticity is not compound specific, for what we still cannot discern if *R. prolixus* is not capable of distinguishing between these two salts, or if they can distinguish, they can learn, but they generalize between salts (*i.e.* distinguishing peripherally between them but transferring what they have learnt for one stimulus to the other).

#### 3.2.4. Chemical pre-exposure to salts

We applied a chemical pre-exposure to NaCl or KCl and analysed then if this treatment had or not an effect over the behaviour of *R. prolixus* confronted to the pre-exposed salt, to the new salt or to both. We assume that the occurrence of a salt-specific effect indicates the existence of a differential detection pathway.

Insects were exposed during 60 minutes to H_2_O, NaCl or KCl. Immediately after their preferences were tested by confronting the same salt or the new one to H_2_O or to each other. Results show that the three chemical pre-exposures (*i.e.* to H_2_O, NaCl and KCl) failed in modulating the innate responses of insects to the salts. Bugs continued to avoid NaCl and KCl presented against H_2_O and evinced no preference when the contingency NaCl/KCl was presented (Fig. 5A, H_2_O/KCl, T = - 2.9, p < 0.01; H_2_O/NaCl, T = −3.8, p < 0.001; KCl/NaCl, T = −0.6, p > 0.05; Fig. 5B, H_2_O/KCl, T = −2.2, p < 0.05; H_2_O/NaCl, T = −2.1, p < 0.05; KCl/NaCl, T = −0.02, p > 0.05; Fig. 5C, H_2_O/KCl, T = −2.5, p < 0.05; H_2_O/NaCl, T = −2.3, p < 0.05; KCl/NaCl, T = −0.7, p > 0.05). These results support the hypothesis that *R. prolixus* cannot distinguish between NaCl and KCl.

**Figure 5.**
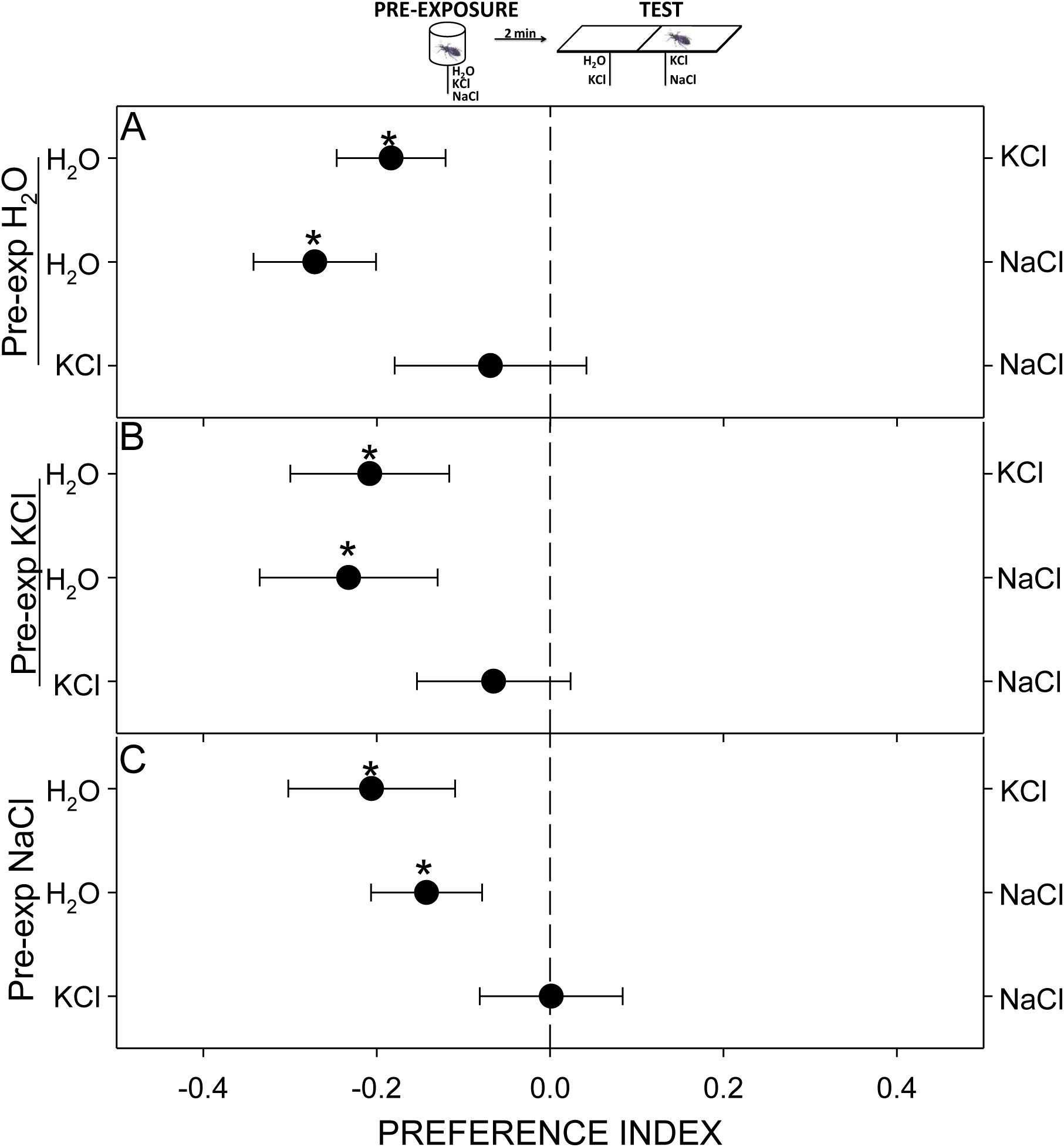
Pre-exposure to salts: effect on the perception of NaCl and KCl. The innate repellence of insects to both salts was not modified by a pre-exposure to H_2_O (A), KCl (B) or NaCl (C). The Preference Index expresses the relative time spent at each side of the arena: 0 = equal time at each side, −1 and 1 = full time spent at the left or right side of the arena, respectively. Each point represents the mean (± s.e.m.) of 30 replicates. Asterisks denote statistical differences (p < 0.05) after a One-Sample T-Test with expected value = 0.

### 3.3. Discrimination between aversive compounds of different gustatory modality, *i.e.* salty and bitter

#### 3.3.1. Innate responses to Caffeine

Results presented in previous sections show that *R. prolixus* avoids walking over an aqueous solution of 1 M NaCl (see Fig. 1). Here we show that an aqueous solution 0.1 M of the alkaloid Caf is also avoided by *R. prolixus* (Fig. 6, H_2_O/Caf, T = −5.5, p < 0.001). However, when this alkaloid is confronted to 1 M NaCl, the insects evinced no preference, remaining the same time at each side of the arena (Fig. 6, Caf/NaCl, T = −0.6, p > 0.05).

**Figure 6.**
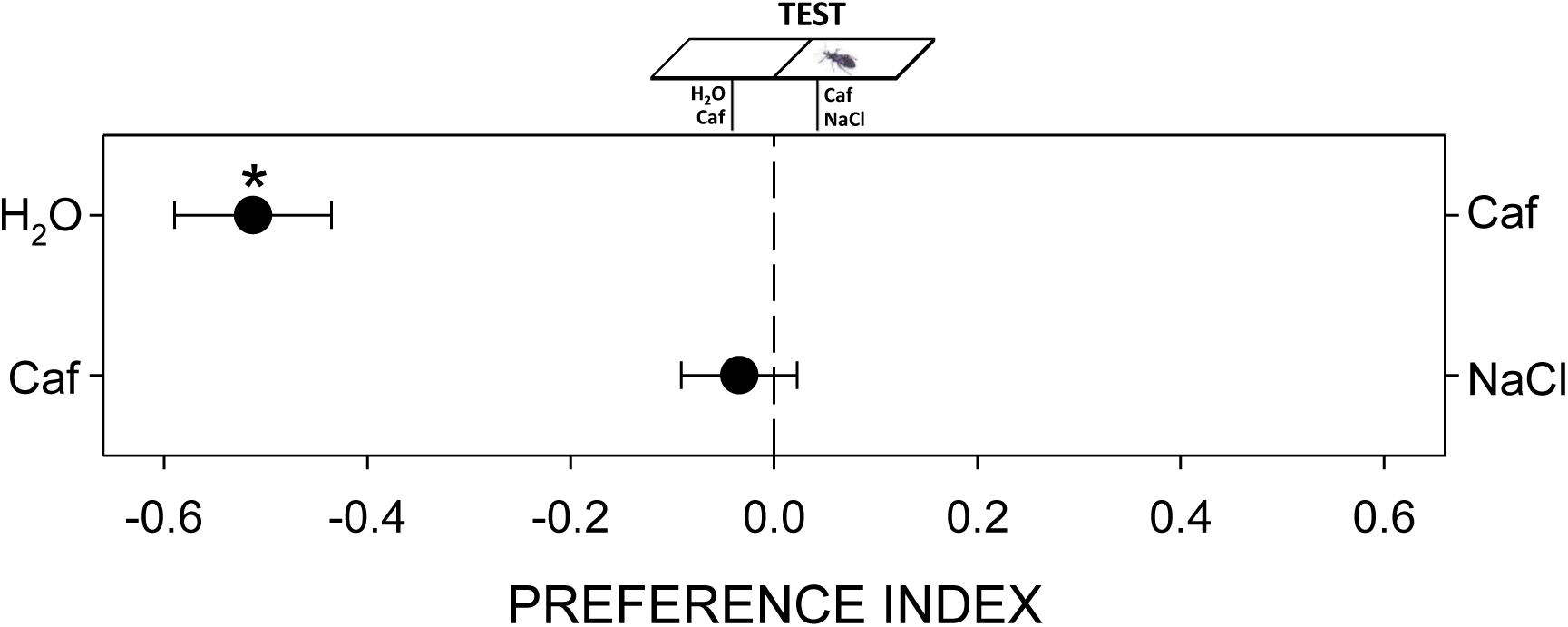
Innate responses to Caf. Insects avoided the caffeine when confronted to H_2_O but exhibited no preference when it was simultaneously presented with NaCl. The Preference Index expresses the relative time spent at each side of the arena: 0 = equal time at each side, −1 and 1 = full time spent at the left or right side of the arena, respectively. Each point represents the mean (± s.e.m.) of 30 replicates. Asterisks denote statistical differences (p < 0.05) after a One-Sample T-Test with expected value = 0.

These results evince that *R. prolixus* can sense the presence of caffeine on the walking substrate, and that this perception generates an avoidance, which is similar in intensity to that exerted by NaCl. Consequently, the question arises again about the capacity of *R. prolixus* to distinguish between these two aversive compounds of radically different chemical identity, *i.e.* a salt and an alkaloid. Next experimental series involving associative and non-associative discriminatory learning protocols were designed to answer this question.

#### 3.3.2. Aversive associative conditioning with salts and alkaloids

For these experimental series we applied a similar operant aversive protocol to that described in section 3.2.3, but pairing in this case the negative reinforcement with the salt, *i.e.* NaCl(-), or with the bitter compound, *i.e.* Caf(-). In both training periods the mechanical disturbance was effective causing insects to avoid the punished side (Fig. 7A, triangle Caf/NaCl(-), T = 8.4, p < 0.001; Fig. 7B, triangle Caf(-)/NaCl, T = −6.6, p < 0.001). However, during subsequent tests insects did not evince any change in their innate behaviour, *i.e.* they showed no preferences (Fig. 7A, circle Caf/NaCl, T = 1.0, p > 0.05; Fig. 7B, circle Caf/NaCl, T = −1.0, p > 0.05).

**Figure 7.**
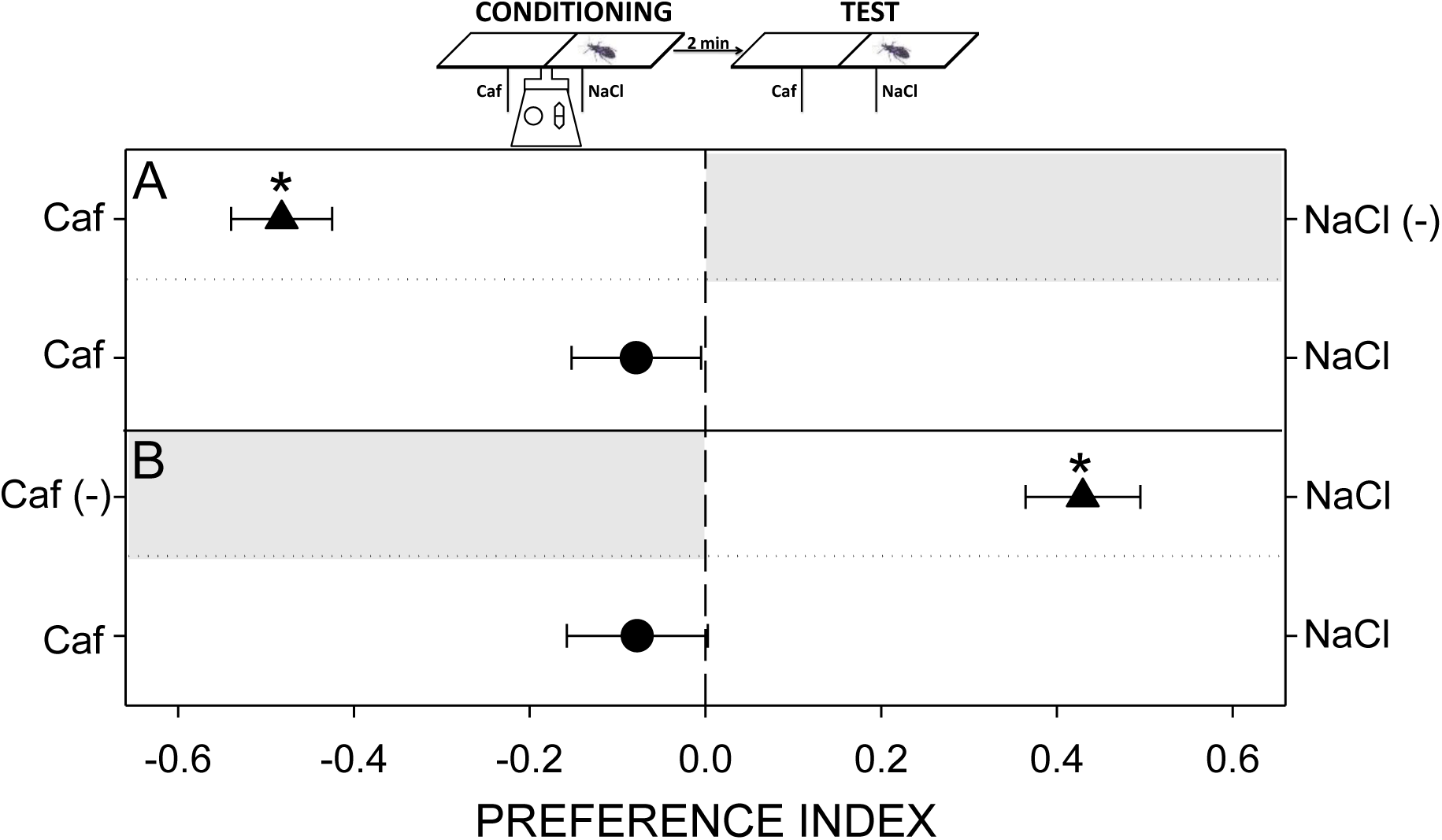
Discriminative assays between aversive stimuli of different gustatory modality through an operant conditioning protocol with NaCl(-) or Caf(-). During trainings (triangles in A and B) insects avoided the punished side of the arena (grey shadowed) regardless if it was loaded with NaCl or Caf. During tests (circles in A and B), no modifications of the innate behaviour were evinced. The Preference Index expresses the relative time spent at each side of the arena: 0 = equal time at each side, −1 and 1 = full time spent at the left or right side of the arena, respectively. Each point represents the mean (± s.e.m.) of 30 replicates. Asterisks denote statistical differences (p < 0.05) after a One-Sample T-Test with expected value = 0.

Even knowing from previous series that our conditioning protocol is effective to generate a spatial memory in these insects, these results do not allow us to determine if *R. prolixus* is not able to distinguish between NaCl and Caf, or if they can do it but they generalize what they have learnt for one compound to the other. In the next series we deal with this question by applying a chemical pre-exposure to each compound and analysing its specificity.

#### 3.3.3. Chemical pre-exposure to salts and alkaloids

Following the 60 minutes chemical pre-exposure to H_2_O, NaCl or Caf, the chemical preferences of insects were tested. No effects were evinced after pre-exposing to H_2_O (Fig. 8A, H_2_O/NaCl, T = −3.8, p < 0.001; H_2_O/Caf, T = −5.6, p < 0.001; Caf/NaCl, T = 0.1, p > 0.05) or NaCl (Fig. 8B, H_2_O/NaCl, T= −2.2, p<0.05; H_2_O/Caf, T= −3.4, p < 0.01; Caf/NaCl, T= 0.7, p > 0.05). In both cases the avoidance of NaCl and Caf when confronted to water, and the lack of preference when the salt and the alkaloid were presented simultaneously were similar to those exhibited by naïve insects (see Figs. 1, 6). However, when insects were pre-exposed to Caf, the aversive response to Caf evanished (Fig. 8C, H_2_O/Caf, T = 1.0, p > 0.05) but the avoidance of NaCl remained intact, effect evinced when confronted to H_2_O or to Caf (H_2_O/NaCl, T = −4.3, p < 0.001; Caf/NaCl, T = 3.6, p < 0.01).

**Figure 8.**
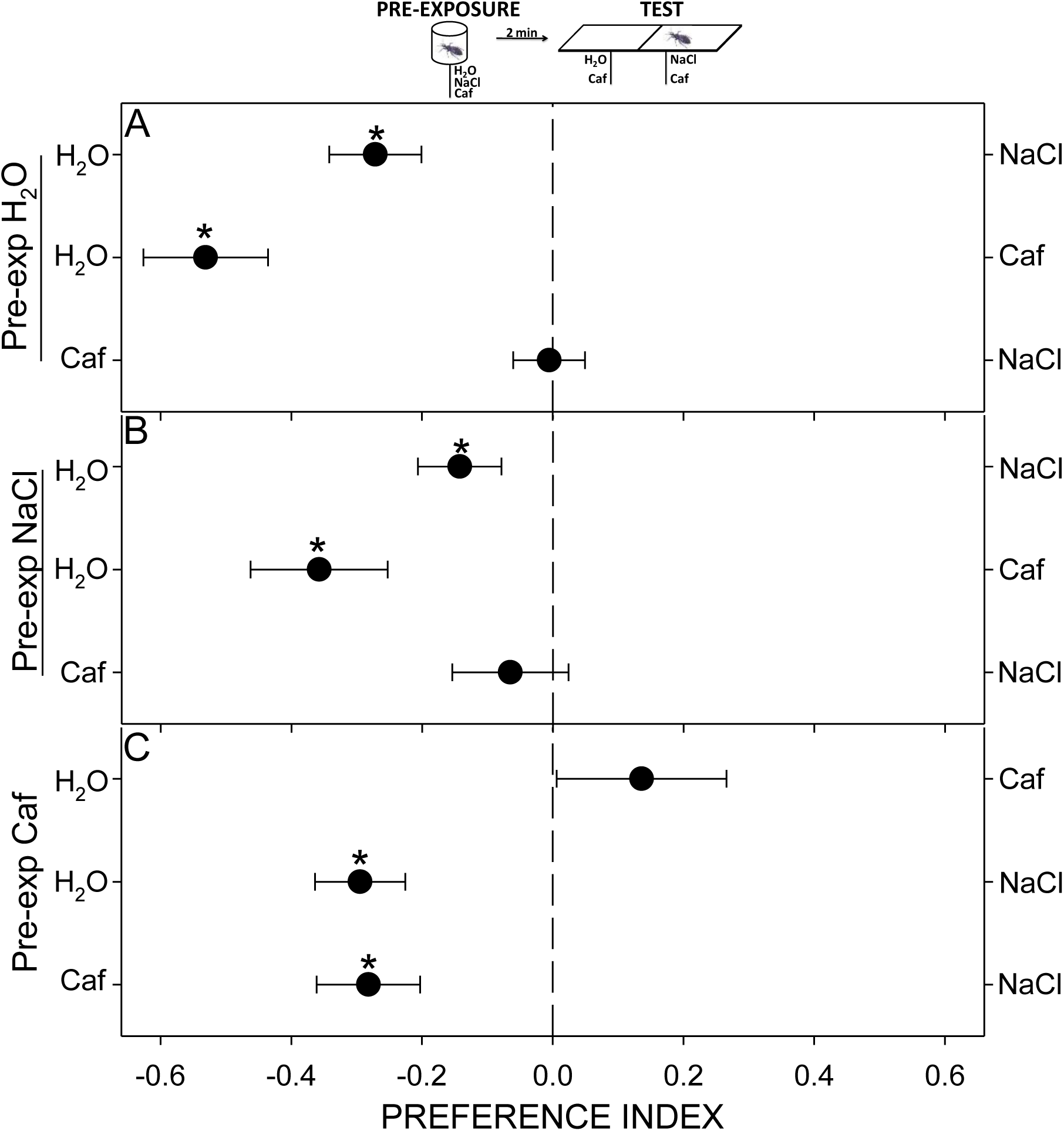
Discriminative assays between aversive stimuli of different gustatory modality through a pre-exposure to NaCl or Caf. A pre-exposure to H_2_O (A) or NaCl (B) did not modify the innate repellence of insects to the salt nor to the alkaloid. However, a pre-exposure to Caf (C) interfered specifically in the perception of the alkaloid but not of the salt. The Preference Index expresses the relative time spent at each side of the arena: 0 = equal time at each side, −1 and 1 = full time spent at the left or right side of the arena, respectively. Each point represents the mean (± s.e.m.) of 30 replicates. Asterisks denote statistical differences (p < 0.05) after a One-Sample T-Test with expected value = 0.

These results, in which the effect of a pre-exposure to Caf is compound-specific, demonstrate that *R. prolixus* can distinguish between the salt NaCl and the alkaloid Caf, and suggest the existence of two different sensory pathways involved in the detection of aversive compounds of different gustatory modality. Additionally, we can validate our method to analyse the taste discrimination in *R. prolixus*, enforcing the evidences presented before in which we suggest that NaCl and KCl are not distinguished by these insects.

### 3.4. Yoke control series

When applying operant conditioning protocols, unpaired yoke controls assure that the experience dependent modulation observed during conditioning is an associative process and not a merely effect of one of the stimuli presented alone. In our experiments, yoke insects were submitted to a pseudoconditioning in which the occurrence of the mechanical punishment was delivered independently from the insect’s chemical choice. As expected, these insects showed a random distribution during trainings regardless the stimulus added at each side of the arena (Fig. S1A, triangles KCl(-)/NaCl(-), T = −0.01, p > 0.05; KCl(-)/NaCl(-), T = −1.0, p > 0.05; Fig. S1B, triangles H_2_O(-)/NaCl(-), T = −0.01, p > 0.05; H_2_O(-)/KCl(-), T = 0.8, p > 0.05; Fig. S1C, triangles Caf(-)/NaCl(-), T = 1.0, p > 0.05; Caf(-)/NaCl(-), T = 1.0, p > 0.05).

The lack of preference when the aversive compounds were confronted to water could be generated by the un-associated delivery of the mechanical disturbance, which could be interfering with the innate chemical preferences of these bugs. As in previous sections, it seems that the negative reinforcement is somehow stronger than the chemical preferences.

During subsequent tests, the animals evinced a behaviour similar to that expressed by naïve ones, distributing randomly over the arena when two aversive compounds were presented in simultaneous (Fig. S1A, B, circles KCl/NaCl, p > 0.05 in 4 cases; Fig. S1C, circles Caf/NaCl, p > 0.05 in 2 cases), and avoiding the aversive stimulus (*i.e.* NaCl and KCl) when confronted to H_2_O (Fig. S1B, circles H_2_O/NaCl, p < 0.05 in 2 cases; H_2_O/KCl, p < 0.05 in 2 cases).

The results of yoke controls confirm that in conditioning groups showed before (see sections 3.2.3 and 3.3.2) the modulation of the behaviour expressed in the tests can be explained by an association between the chemical stimulus and the negative reinforcement and not by a non-associative modulation caused by the chemical exposure or the mechanical disturbance alone.

## 4. Discussion

In this work we examined the capacity of *R. prolixus* to perceive aversive chemical compounds and to distinguish between stimuli of the same or of different gustatory modalities, *i.e.* salty (NaCl and KCl) and bitter (Caf). Results here presented strongly suggest that *R. prolixus* is not able to distinguish between the two salts: NaCl and KCl. Conversely, we demonstrate that these insects can distinguish between salts and alkaloids, probably by detecting these compounds of different gustatory modality via different gustatory receptors. Still, electrophysiological recordings would be needed to confirm our findings.

Differently from olfactory sensory neurons, gustatory neurons of insects can bear different types of receptors in the surface of the same dendrite. As a result, different molecules (*e.g.* “A” and “B”) reaching different receptors of the same neuron can generate the same electric transduction to higher order brain structures, causing a deficit in the capacity to distinguish between A and B. This apparent disadvantage may in fact be an adaptive value, as animals might only need to detect diverse compounds as a unique stimulus signalizing a potential harm and categorize them as “aversive” independently from their chemical identity. In an extreme case, all aversive chemical stimuli could be perceived equally as negative inputs, and evoke the same escape/reject response.

There are many reports presenting evidence that mammals can distinguish between certain salts (Spector *et al*., 1996). In our work we show that *R. prolixus* is not able to differentiate between NaCl and KCl, but it could discriminate between NaCl and a bitter alkaloid such as the caffeine. These results confirm that, as it happens in most animals, it is quite difficult for *R. prolixus* to distinguish between different compounds of the same gustatory modality (*e.g.* salty) and more feasible to distinguish between tastants of different modalities (*e.g.* salty and bitter).

Commercially available repellents for haematophagous insects typically make use of the olfactory sense of these insects to keep them away from our skin. However, the gustatory sense is by far less taken into consideration for the chemical formulation of repellents. We show here the existence of non-toxic contact repellents (as salts and bitter compounds) that could be used simultaneously with olfactory stimuli as to obtain a multimodal aversive product. Odours acting at a longer distance and tastants at a shorter range could significantly increase the repellency and eventually the biting deterrence efficiency of a final product. Moreover, being barely volatile compounds, they could probably present a higher effective action along time than odours. Still, experiments combining stimuli from both sensory modalities would be needed to confirm this suggestion.

Learning and memory has been shown to occur in most animals. The importance of learning and memory in haematophagous insects has been suggested already more than 60 years ago, when Nielsen and Nielsen (1953) described that mosquitoes return to places where they had previously fed, a behaviour known as homing behaviour. Other authors also suggested the existence of a spatial memory in mosquitoes, as they reported that in natural environments they are capable of associating particular odours of the environment with favourable oviposition sites or hosts (Alonso *et al*., 2003; Chilaka *et al*., 2012; Kaur *et al*., 2003; McCall and Eaton, 2001; McCall *et al*., 2001; McCall and Kelly, 2002; Menda *et al*., 2012; Mwandawiro *et al*., 2000; Sanford and Tomberlin, 2011; Tomberlin *et al*., 2006; Vinauger *et al*., 2014). Alonso and Schuck-Paim (2006) discuss about particular cases found in the literature in which authors erroneously ascertain the existence of learning and memory processes in haematophagous insects. Many advances have been made in understanding the cognitive abilities of haematophagous insects other than mosquitoes in the last decade. For example, in a controlled experimental setup, the kissing bug *R. prolixus* started to walk towards or against an originally neutral odour after an appetitive or aversive conditioning, respectively (Vinauger *et al*., 2011a, b). Same authors found that even if *R. prolixus* did not present a preference when odours from a live rat or quail were presented simultaneously, an aversive conditioning generated an aversion to odours from the punished host (Vinauger *et al*., 2012). Taking advantage of the proboscis extension response elicited by triatomine insects confronted to a warm surface, Vinauger and collaborators (2013) demonstrated that the PER of *R. prolixus* can be modulated by non-associative and associative learning forms. In a completely different context, it was shown that the escape response of *R. prolixus* to the alarm pheromone can be widely modulated by associative and non-associative conditioning protocols (Minoli *et al*., 2013). In a feeding context, it was reported that the addition of bitter compounds to an appetitive diet inhibits the feeding behaviour of *R. prolixus* (Pontes *et al*., 2014). Same authors also found that a brief pre-exposure to bitter compounds negatively modulates the motivation of bugs to feed on an appetitive solution. Additionally, it was shown that triatomines’ cognitive abilities follow a circadian rhythm, performing well during the night, but not during the day (Vinauger and Lazzari, 2015). As well, studying the repellent effect of new non-toxic molecules for *R. prolixus*, Asparch and collaborators (2016) found that bugs are innately repelled by different bitter molecules, and that this repellence can be modulated by associative and non-associative forms of learning. Indeed, after an aversive operant conditioning, the behaviour of *R. prolixus* changed from avoidance to indifference or even to preference, according with the protocol applied (Asparch *et al*., 2016). In a completely different behavioural context, Mengoni and collaborators (2017) showed that the innate attractive response to faeces of the kissing bug *Triatoma infestans* can be switched to avoidance if an aversive conditioning protocol is applied. Finally, Minoli and collaborators (2018) were able to demonstrate that the learning efficiency of *R. prolixus* under an aversive operant conditioning tested over a spatial preference walking arena, is highly dependent on the sensory modality of the conditioned stimulus.

This work enriches the knowledge about gustatory perception in an haematophagous vector insect, and highlights the utilization of the learning process as an indispensable tool for the determination of their discrimination capacities. Moreover, taking in consideration that *R. prolixus* is an insect-vector of a human disease and that its genome has been recently sequenced (Mezquita *et al*., 2015), it can become a promising model in the learning and memory field. Knowledge about the gustatory system of vector insects and its inherent plasticity provides opportunities to develop new sustainable methods to reduce or even prevent vector-host contacts and disease transmission.

## 5. Symbols and abbreviations

GR: Gustatory receptor
GRN: Gustatory receptor neuron
H_2_O: Distilled water
NaCl: Sodium chloride
KCl: Potassium chloride
Caf: Caffeine anhydrous
12: 12 h. L / D: 12 hours of light and 12 hours of darkness
PI: Preference index
ENaC: Epithelial sodium channels
(-): sign added besides the stimulus punished during training

## 6. Acknowledgements

Authors thank to G. P. Jerez Ferreyra for his invaluable help rearing and maintaining the insect colony.

## 7. Author contributions

S.Ma. performed experiments, analysed data and wrote manuscript.

A.C performed experiments.

Y.A. performed experiments.

R.B. designed experiments, wrote manuscript.

S.Mi. designed experiments, analysed data and wrote manuscript.

---

**Figure S1.**
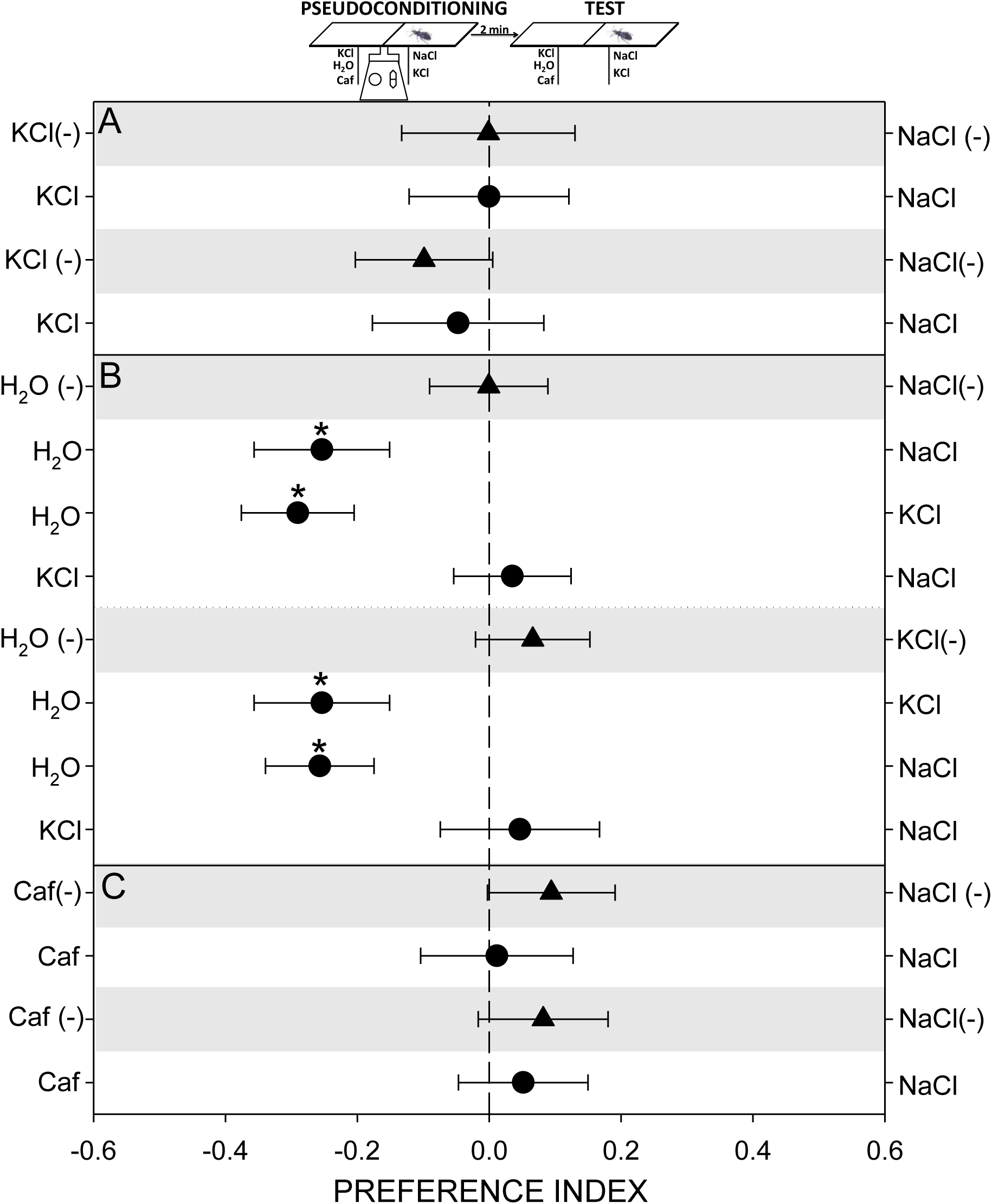
Operant yoke control experiments: unpaired presentation of the chemical stimulus and the mechanical disturbance. During trainings (triangles in A, B and C), no chemical preferences were observed (grey shadows evince that mechanical punishment could be delivered at both sides of the arena). During tests (circles in A, B and C), no modifications of the innate behaviours were displayed. (A) Yoke control of Fig. 3. (B) Yoke control of Fig. 4. (C) Yoke control of Fig. 7. The Preference Index expresses the relative time spent at each side of the arena: 0 = equal time at each side, −1 and 1 = full time spent at the left or right side of the arena, respectively. Each point represents the mean (± s.e.m.) of 30 replicates. Asterisks denote statistical differences (p < 0.05) after a One-Sample T-Test with expected value = 0.

